# A novel mode of photoprotection mediated by a cysteine residue in the chlorophyll protein IsiA

**DOI:** 10.1101/2020.12.24.424336

**Authors:** Hui-Yuan Steven Chen, Dariusz M. Niedzwiedzki, Anindita Bandyopadhyay, Sandeep Biswas, Himadri B. Pakrasi

**Author notes:** Address correspondence to Himadri B. Pakrasi, e. Bayer Crop Science, Chesterfield, MO 63017, USA.

## Abstract

Oxygenic photosynthetic organisms have evolved multitude mechanisms for protection against high light stress. IsiA, a chlorophyll *a*-binding cyanobacterial protein serves as an accessory antenna complex for photosystem I. Intriguingly, IsiA can also function as an independent pigment protein complex in the thylakoid membrane and facilitate dissipation of excess energy, providing photoprotection. The molecular basis of IsiA-mediated excitation quenching mechanism remains poorly understood. In this study, we demonstrate that IsiA uses a novel cysteine-mediated process to quench excitation energy. The single cysteine in IsiA in the cyanobacterium *Synechocystis* 6803 was converted to a valine. Ultrafast fluorescence spectroscopic analysis showed that this single change abolishes the excitation energy quenching ability of IsiA, thus providing direct evidence of the crucial role of this cysteine residue in the energy dissipation from excited chlorophylls. Under stress condition, the mutant cells exhibited enhanced light sensitivity, indicating that the cysteine-mediated quenching process is critically important for photoprotection.

## Introduction

Exposure of cyanobacteria, algae and green plants to high light intensities often leads to damage to their photosynthetic apparatus. Cyanobacteria are oxygenic photosynthetic prokaryotes that depend on sunlight for their growth and survival. During billions of years of their evolution, these microbes have developed a number of strategies to modulate light absorption and dissipation to ensure maximal photosynthetic productivity and minimal photodamage to cells under extreme light and limiting nutrient conditions. Iron deficiency is a common nutrient stress in various cyanobacterial habitats^2-6^. In cyanobacteria, iron is mostly used in photosynthetic reaction center complexes and in iron-depleted environments, their photosynthetic machinery exhibits certain adaptive changes such as decreased amount of chlorophyll (Chl)-binding proteins and phycobilisomes^7-9^. Another adaptive photoprotective strategy that cyanobacteria have evolved is the induction of the iron <u>s</u>tress-induced protein <u>A</u> (IsiA)^8,10^.

IsiA is a Chl *a*-binding membrane protein which was first discovered in cyanobacteria grown in iron-free media^8,10^. Later studies showed that IsiA can also be induced by oxidative stress, high salt, heat stress, and in particular, high light^11-16^. IsiA belongs to a superfamily of antenna proteins with six transmembrane helices^17^, and is highly homologous with CP43, an intrinsic antenna protein of photosystem II (PSII). However, unlike CP43, IsiA is mainly found to be associated with photosystem I (PSI), and forms PSI_3_-IsiA_18_ supercomplexes^18-20^. Time-resolved spectroscopic studies showed that the energy transfer from IsiA to PSI and between IsiA copies in PSI-IsiA supercomplexes is fast and efficient ^21,22^. Since one IsiA binds 17 Chl *a* molecules^20,22,23^ and one PSI monomer binds 96 Chl *a*^24^ molecules, the outer IsiA ring can theoretically increase the absorption cross-section of the PSI_3_-IsiA_18_ supercomplex by at least 81% compared to a PSI trimer alone. *In vivo*, IsiA increases the effective absorption cross-section of PSI by ∼60%^25^. These results showed that in the PSI_3_-IsiA_18_ supercomplex, IsiA serves as an accessory antenna for PSI.

In addition to the PSI_3_-IsiA_18_ supercomplex, IsiA-only pigment protein complexes have been observed in cyanobacterial cells after prolonged growth under iron-depleted conditions^26^ as well as under high light conditions^16^. Such IsiA-only complexes have been suggested to be involved in non-photochemical quenching processes^27^. Previous studies showed that an *isiA* deletion strain is more light-sensitive, suggesting that IsiA plays a significant role in providing photoprotection^13,27,28^. Using time-resolved fluorescence spectroscopy, it has been determined that the accumulation of IsiA-only complexes in the cells results in strong quenching of IsiA fluorescence, suggesting its photoprotective role^28,29^. However, the mechanism of IsiA-mediated excitation quenching is not fully understood. Earlier, it was proposed that the carotenoids present in the IsiA protein are solely responsible for the quenching of the singlet state of Chl *a* ^30,31^. However, this premise has recently been questioned and instead, we have proposed a novel cysteine-mediated quenching mechanism for this cyanobacterial pigment protein^32^. Such a mechanism was originally proposed for the Fenna-Matthews-Olson (FMO) protein complex in anoxygenic green sulfur bacteria^33^.

A recent single-particle cryo-electron microscopy study reported the structure of the PSI_3_-IsiA_18_ supercomplex from *Synechocystis* sp. PCC 6803 (hereafter *Synechocystis*) at a 3.5 Å resolution^34^. This pioneering study has provided significant insights into the protein-protein interactions in the PSI-IsiA supercomplex and identified the potential terminal emitters, the Chl *a* molecules located at the PSI-IsiA interface, in IsiA^20^. Since then, two other studies have elucidated the structures of IsiA in PSI-IsiA complexes from two other cyanobacterial species at even higher resolutions^35,36^. Although high resolution structure of an IsiA-only complex is yet to be determined, the available structures of IsiA in the PSI-IsiA supercomplex help shed new lights into the excitation energy quenching process in this protein. *Synechocystis* IsiA has a unique cysteine, C260, which is part of an ‘AYFCAVN’ motif^37^ located at the luminal side of the transmembrane helix V^20^. An alignment of the amino acid sequence of IsiA from several representative cyanobacterial strains shows that this motif is highly conserved in cyanobacteria (Figure 1). Interestingly, in CP43, a protein that does not quench excitation on its own, the cysteine residue in this conserved motif is substituted by valine. According to the proposed cysteine-mediated quenching mechanism^32^, a valine at that position would be unable to facilitate the quenching process. In the current study, we performed site-directed mutagenesis to construct *Synechocystis* strains in which the unique cysteine in IsiA was replaced with a valine. Remarkably, with this single amino acid change, the mutant IsiA was unable to quench excitation energy, but was still capable of serving as an efficient light-harvesting antenna for PSI. Furthermore, the C260V mutant strain was more light-sensitive under stringent iron-depleted conditions but had a faster growth rate compared to the wild type (WT) cells in iron-replete conditions under high light.

**Figure 1.**
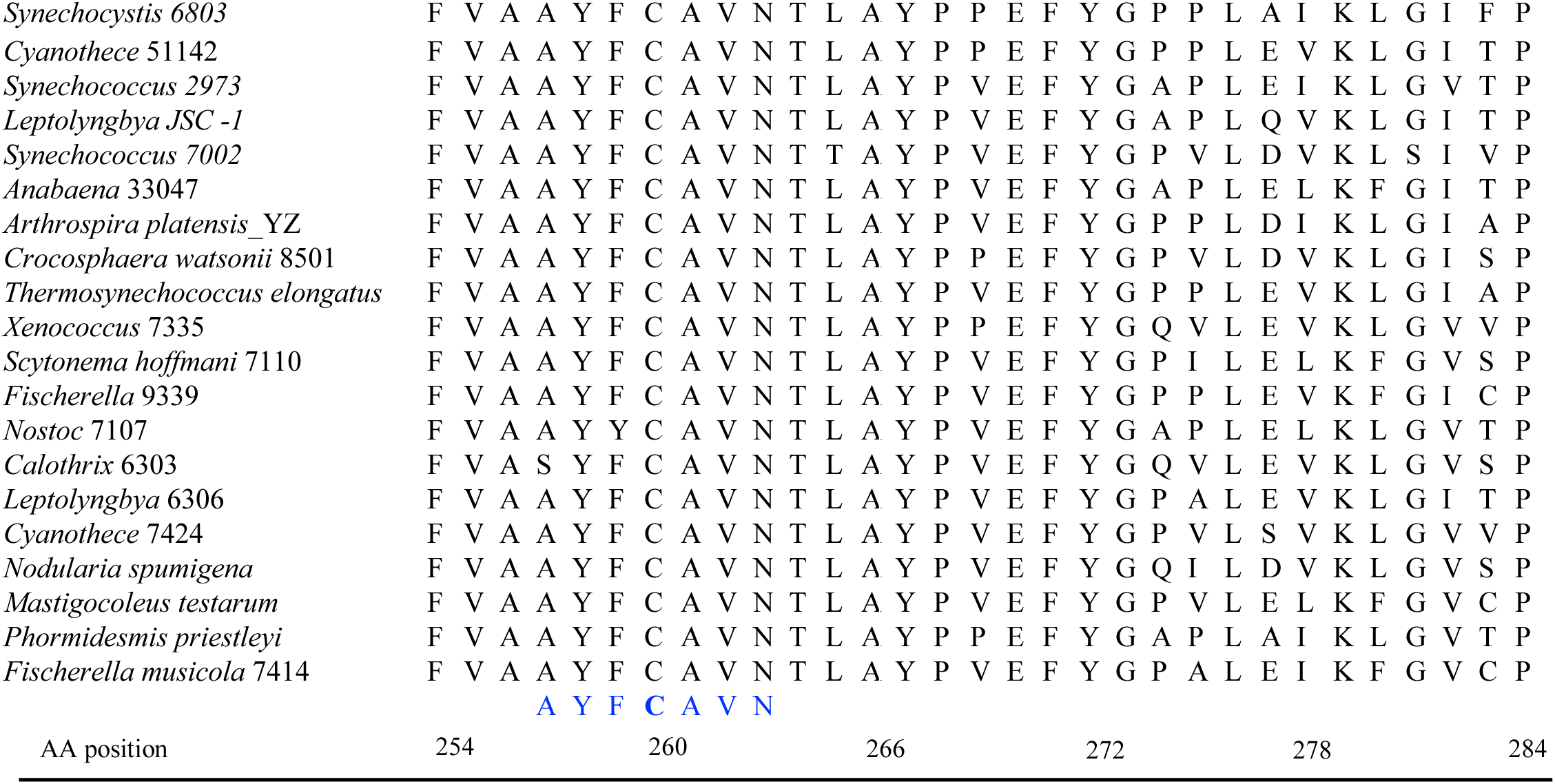
Sequence alignment of the IsiA protein showing the conserved cysteine residue. Sequence alignment of the IsiA protein from 24 strains representative of unicellular, filamentous, diazotrophic and non-diazotrophic cyanobacteria. The sequences were aligned with ClustalW within MEGA 7^1^. The cysteine residue in the ‘AYFCAVN’ motif is conserved across the cyanobacterial strains

## Results

### Construction of C260V and C260V-His *Synechocystis* strains

The C260V mutation was introduced into the WT *Synechocystis* strain using a CRISPR/Cas12a (Cpf1) system^38^. The resulting C260V strain is a markerless mutant with the cysteine to valine substitution as the only change. All the physiological comparisons in this study were done using this mutant and WT *Synechocystis* strains. On the other hand, for the biophysical and biochemical studies, pure IsiA and PSI-IsiA supercomplexes were needed. In our previous study, a histidine tagged IsiA transgenic line was generated which enabled us to purify the individual supercomplexes^32^. In this study, the C260V mutation was introduced into this IsiA-His strain via double homologous recombination. The resulting strain, C260V-His, was grown in iron-depleted conditions that induce *isiA* expression. The mutant IsiA and PSI-IsiA supercomplexes were purified from the C260V-His strain by affinity chromatography followed by rate-zonal centrifugation.

### Biochemical and spectroscopic analysis of mutant PSI-IsiA and IsiA protein complexes

To assess how the single amino acid substitution, C260V, affects the biophysical properties of the mutant PSI-IsiA and IsiA aggregates, pure IsiA and PSI-IsiA supercomplexes were isolated. Figure 2A shows the purified protein complexes. Band 1 (top) and band 4 (bottom) were analyzed by immunoblotting. The proteins in both bands were fractionated by SDS-PAGE and probed using antisera raised specifically against PsaA and IsiA (Figure 2B). The results showed that band 1 contained the IsiA-only complex without any PSI contamination whereas band 4 contained PSI-IsiA supercomplexes. These samples will be abbreviated as C260V IsiA and PSI-C260V IsiA, respectively. Sample purity was also confirmed by analysis of room temperature absorption spectra of both preparations (Figure 2C, D). Comparison of the spectroscopic profiles of WT-IsiA and C260V-IsiA shows that both complexes have essentially identical Chl *a* Q_y_ band with an absorption maximum at 670.8 nm (Figure 2C) indicating no PSI contamination, that is known to shift position of the Chl *a* Q_y_ band by a few nanometers toward longer wavelengths^32,39,40^. Such a shift was observed in the spectra of both WT and mutant PSI-IsiA complexes (673.8 nm, Figure 2D). In addition, the higher absorbance in the 450 nm to 500 nm region indicated an enhanced level of carotenoids in the C260V IsiA sample. This may be a physiological response to the mutation.

**Figure 2.**
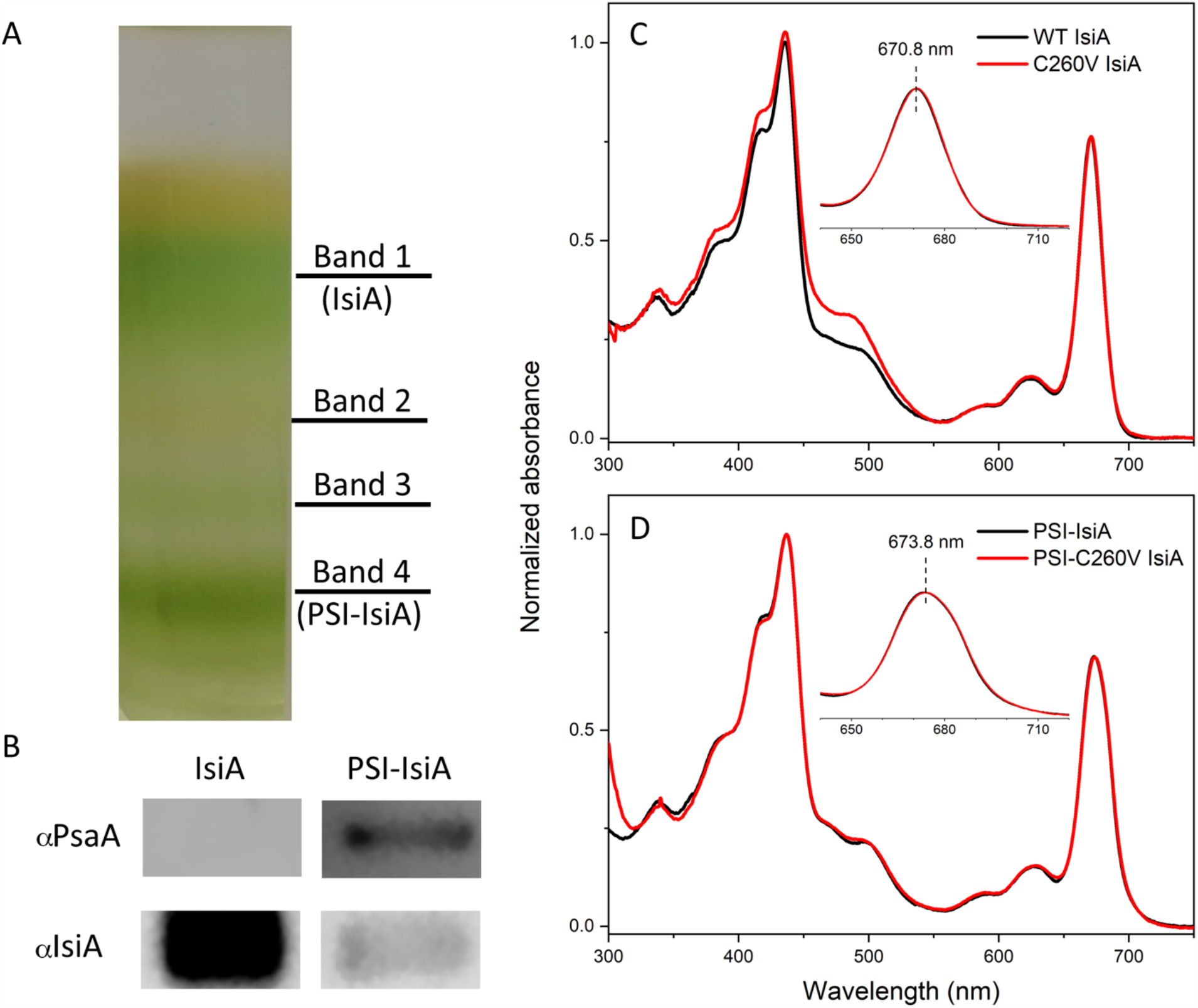
Purification of C260V IsiA and PSI-C260V IsiA pigment protein complexes from the C260V-His tagged strain and their basic spectroscopic characterization. (A) Pigment-protein bands obtained from sucrose gradient ultracentrifugation with the IsiA and PSI-IsiA bands labeled; (B) analysis of IsiA and PSI-IsiA sample purity by immunoblotting using antisera raised against PsaA (αPsaA) and IsiA (αIsiA), respectively; Room temperature absorption spectra of (C) WT and C260V IsiA and (D) isolated WT and C260V PSI-IsiA complexes.

### Chl *a* fluorescence decay dynamics in WT and C260V IsiA complexes

Our previous studies of Chl *a* fluorescence decay in the WT IsiA demonstrated that Chl *a* fluorescence lifetime is sensitive to the presence of the reducing agent, sodium dithionite, in the sample buffer ^32^. Therefore, we hypothesized that Chl *a* lifetime extension is caused by an inhibition of the cysteine-mediated excitation quenching and reasoned that without the unique cysteine residue, this quenching mechanism in IsiA would be significantly affected or even completely absent, giving rise to longer Chl *a* fluorescence decay in the C260V IsiA mutant. This Hypothesis was verified as demonstrated in Fig. 3. Our comparison of Chl *a* fluorescence decay in WT and C260V IsiA samples (Figure 3) showed that the fluorescence lifetime of the mutant IsiA is even longer than that of the chemically reduced WT IsiA, indicating the absence of excitation energy quenching in the mutant strain.

**Figure 3.**
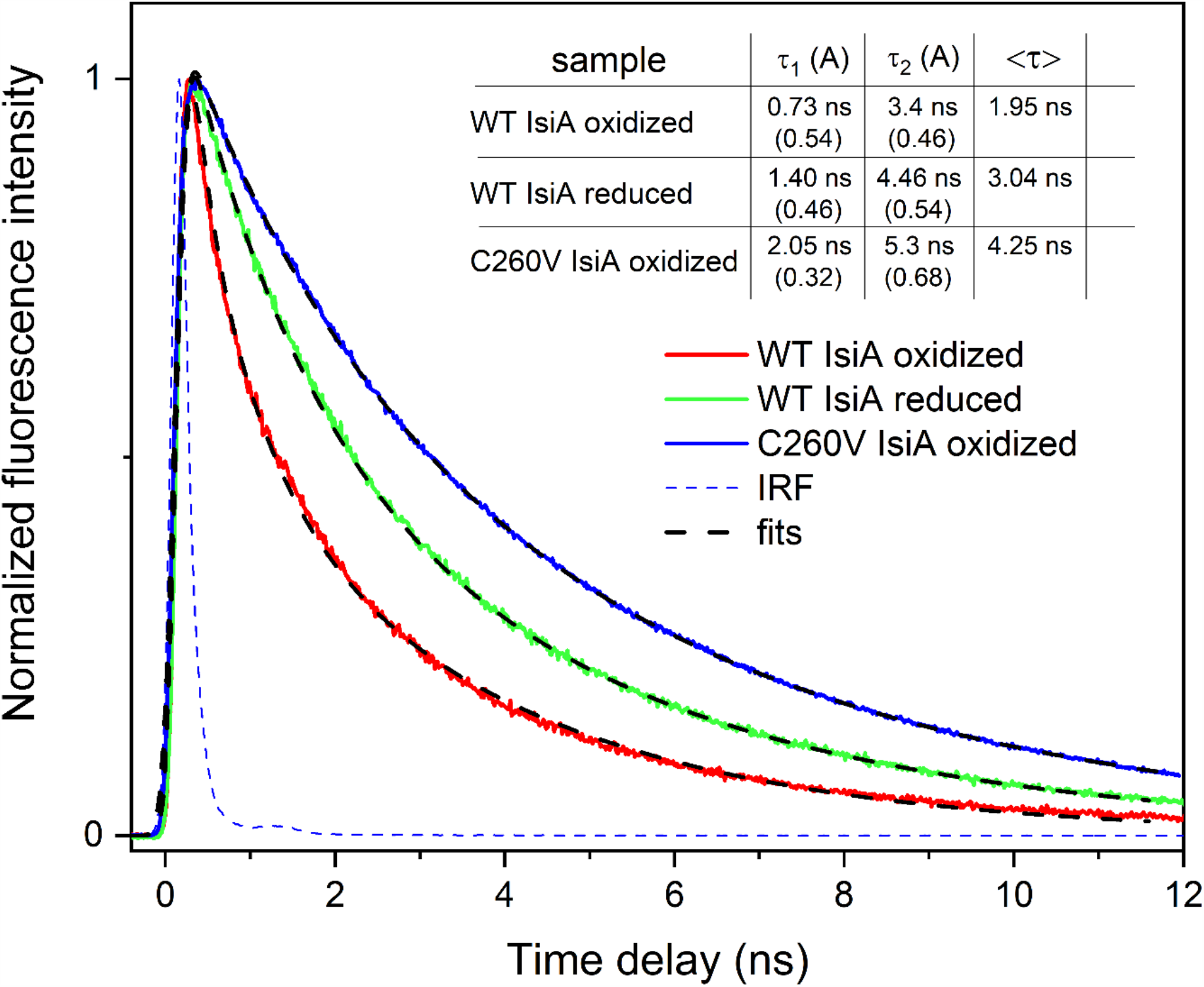
Fluorescence decay dynamics of IsiA-bound Chl *a* in WT and C260V strains, under oxidative (ox, buffer as is) and reducing (red, after addition of 10 mM sodium dithionite) conditions. Fluorescence decay was recorded at 684 nm at room temperature. IRF – instrument response function. The insert table shows fitting results with lifetimes and amplitudes of contributing kinetic components as well amplitude weighted lifetime <τ>. The signals were normalized for better comparability.

Next, time-resolved fluorescence spectra revealed that both WT and C260V IsiA proteins, when assembled into supercomplexes with PSI, show substantial shortening of Chl *a* fluorescence decay, demonstrating that the mutant antenna complex was capable of efficient transfer of excitation energy to PSI (Figure 4). Figures 4A and B show two-dimensional pseudo-color fluorescence decay profiles recorded for both supercomplexes. Cryogenic temperature allows recording of fluorescence from PSI (720 nm band) and therefore, our measurements were performed at 77 K. Figures 4C and D show time-integrated fluorescence spectra that were generated by integration of all of the time-resolved spectra and, in principle, should be equivalent to the steady-state fluorescence spectra of the supercomplexes. Small differences visible in spectral profiles (670-680 nm) and the residual long-lived signal at ∼680 nm in the mutant sample are associated with larger scattering of the excitation beam and possible residual contamination with free Chl *a*. Fluorescence decay traces of Chl *a* associated with IsiA and PSI, normalized to unity at their maxima (Figure 4E), revealed that fluorescence decays of IsiA-bound Chl *a* were similar in both WT and C260V samples. Therefore, both WT and mutant IsiA function equally well as light harvesting antenna and donors of excitation energy to PSI.

**Figure 4.**
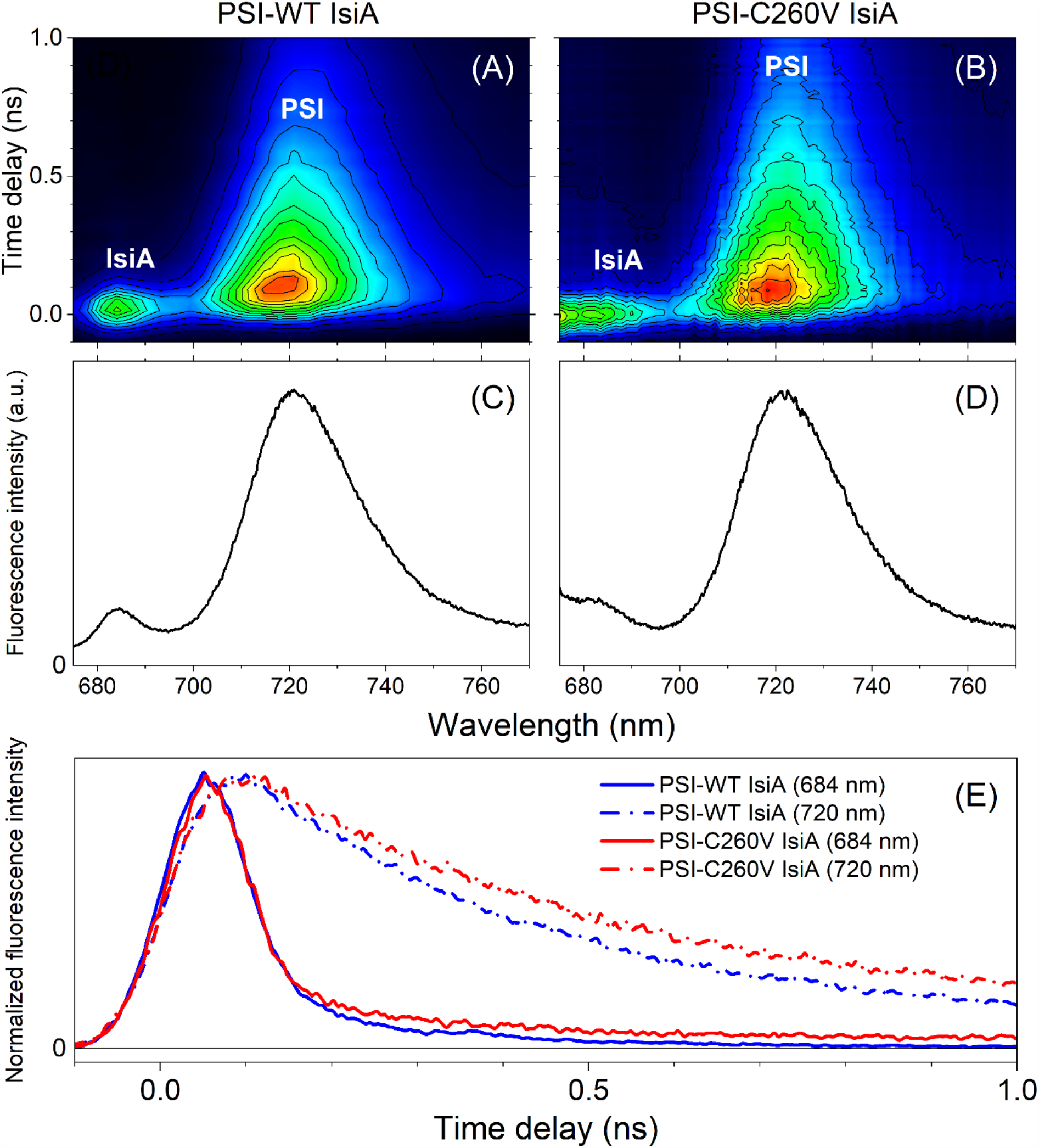
Time-resolved fluorescence from PSI-IsiA supercomplexes at 77 K. (A, B) Two-dimensional, pseudo-color fluorescence decay profiles of PSI-WT and PSI-C260V IsiA supercomplexes, (C, D) Time-integrated spectra that correspond to steady-state fluorescence emission from both supercomplexes. (E) Comparison of IsiA-bound Chl *a* fluorescence decay in both samples. The kinetic traces are normalized to their maxima for better comparability. The samples were excited at 660 nm.

### Changes in composition of pigments and key membrane proteins in the C260V mutant and WT *Synechocystis* strains

To assess the impact of the C260V mutation on cellular physiology, we analyzed pigment and protein compositions of the mutant cells grown under different conditions. The absorption spectra of C260V and WT cultures grown under low light and iron-replete conditions had no noticeable difference (Figure 5A). This was expected because *isiA* is not expressed under these conditions, and, therefore, the mutation is not likely to affect the physiology of the cells. In contrast, when grown under high light and iron-replete conditions, *isiA* expression was induced in both cultures, as indicated by the blue shift of the Q_y_ absorption band (Figure 5B)^37^. Absorption spectra of cells grown under low as well as high light and iron-deplete conditions show that both strains have the blue shift of Chl *a* Q_y_ absorption band from 678 nm to 671 nm. Interestingly, when the C260V mutant is subjected to high light and iron stress, absorption band at 671 nm is significantly reduced, implying a decreased Chl *a* content (Figure 5C). In addition, the Chl *a* Q_y_ band is markedly lower compared to the phycocyanin peak at 625 nm, suggesting an altered pigment composition in the mutant strain.

**Figure 5.**
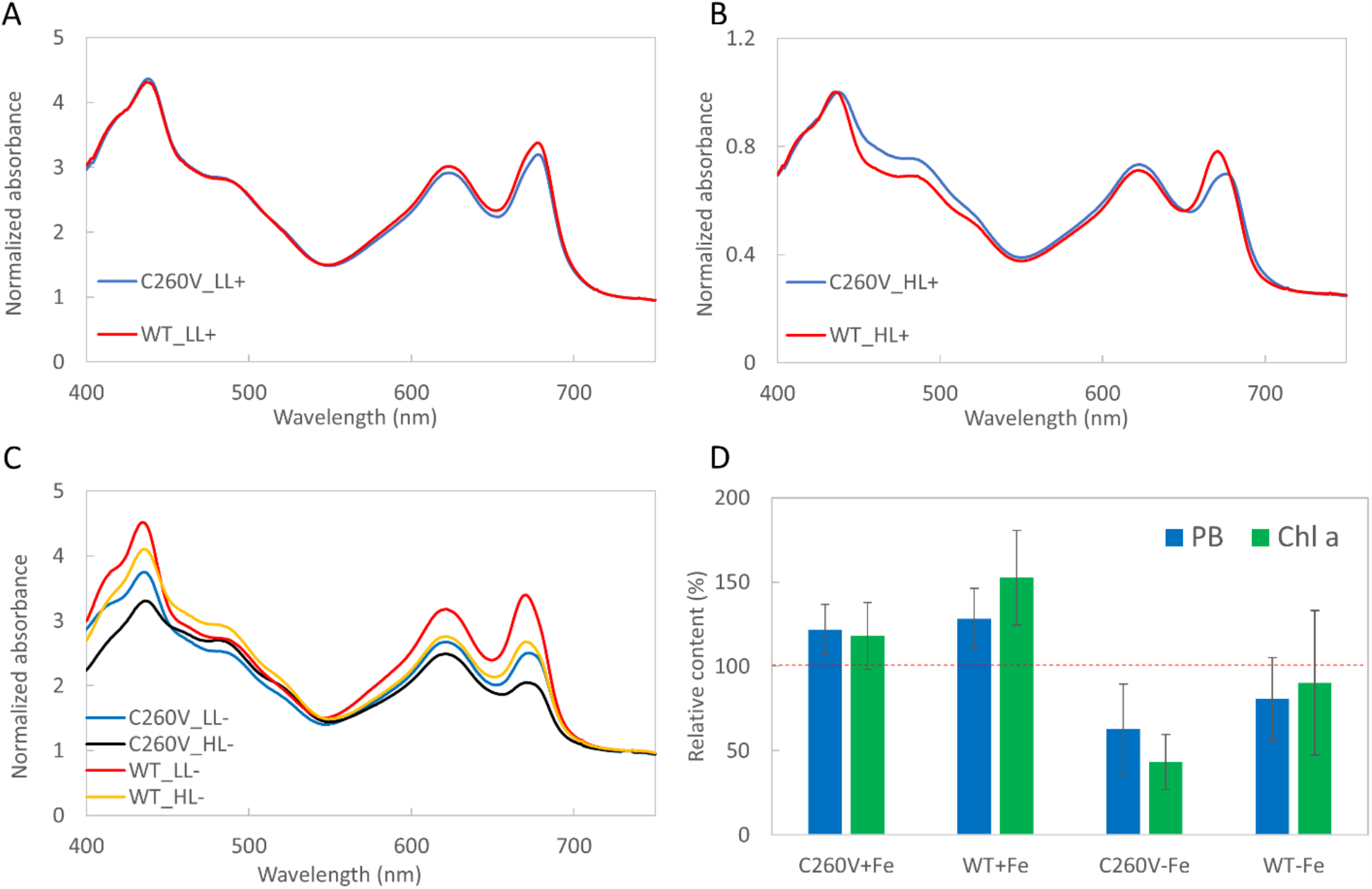
Absorption spectra and relative pigment content of WT and C260V strains. Cultures were grown in multicultivators under (A) 200 μmol photons m^−2^ s^−1^ (low light, LL) with sufficient iron (+), (B) 800 μmol photons m^−2^ s^−1^ (high light, HL) with sufficient iron (+) and (C) under low light (LL) and high light (HL) in the absence of iron (-). (D) Relative phycobilin (PB) and Chl *a* contents per cell in WT and C260V strains under iron-replete and iron-depleted conditions. The spectra were normalized to the absorption at 730 nm. The pigment content of both strains grown under low light is represented as 100% (red dashed line), and the bars represent the phycobilin and Chl *a* content of cultures grown under high light. Error bars represent standard deviation across triplicate biological samples.

To further investigate the impact of high light on the mutant cell physiology, we compared the pigment and protein compositions of the cells grown under high as well as low light intensities (Figure 5D). Under high light and iron-replete conditions, the C260V mutant showed a ∼20% increase in both phycobilin and Chl *a* content, while the WT showed a ∼30% increase in phycobilin and ∼50% increase in Chl *a* content. On the other hand, under high light and iron-depleted conditions, phycobilin and Chl *a* content in both strains decreased pronouncedly. Most striking was the change in the Chl *a* content of the C260V mutant, a ∼55% reduction. The higher phycobilin to Chl *a* ratio in the C260V mutant under high light and iron-depleted conditions (Figure 5C) could be attributed to this severe reduction in Chl *a* content. This led us to investigate which of the Chl *a*-binding proteins were specifically lost to cause such a remarkable decrease in the Chl *a* content.

Under high light and iron-replete conditions, both the C260V mutant and the WT cells showed an increase in PSII and Chl *a* content (Figure 6A). Neither strain expressed *isiA* when grown under low light in the presence of iron and hence the relative IsiA content under these conditions is not shown (Figure 6A). In contrast, when grown under high light, both strains produced IsiA (Figure 6C). However, the high light-induced increase in Chl *a* content was not as marked in C260V mutant as compared to the WT strain, while the C260V mutant exhibited higher PSII content. This is because of the much higher IsiA content in the WT cells. Interestingly, under iron-depleted conditions, the PSI, IsiA and Chl *a* content of the C260V mutant decreased significantly (Figure 6B), whereas the WT cells showed a slight decrease in the PSI and PSII content and a more noticeable decrease in the IsiA content, causing a moderate decrease in the total Chl *a* content (Figure 6B). In addition, the ratio of IsiA content to PSI content in the C260V mutant became much lower as compared to the WT strain, suggesting a significant decrease in the IsiA-only complex in the C260V mutant under high light conditions due to the mutation. These findings indicate that the lack of the quenching ability in the mutant IsiA protein under iron-depleted and high light conditions resulted in severe photodamage of IsiA and PSI.

**Figure 6.**
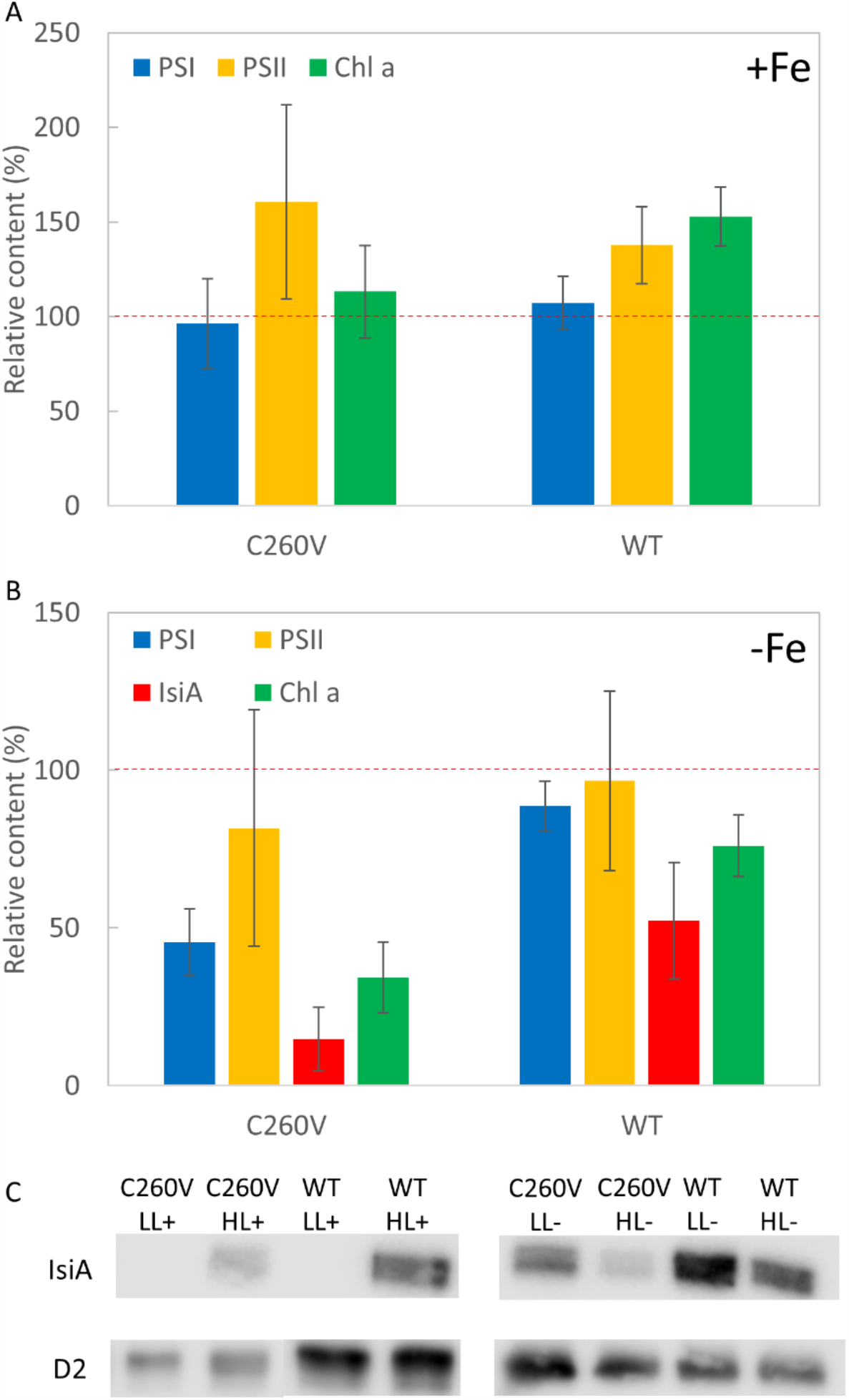
Relative abundance of Chl *a*, IsiA and photosystems of WT and C260V mutant strains. Chl *a*, IsiA, PSI and PSII contents of the C260V and WT cells grown in (A) iron-replete and (B) iron-depleted media. The protein and Chl *a* content of both strains grown under low light (LL) are represented as 100% (red dashed line), and the bars represent the relative protein and Chl *a* content of cells grown under high light (HL). (C) Immunoblot analysis of thylakoid membranes from C260V and WT cells grown under iron-replete conditions and low light (LL+), iron-replete conditions and high light (HL+), iron-depleted conditions under low light (LL-), and iron-depleted conditions under high light (HL-). Samples were probed with specific antisera against IsiA and D2, respectively. Error bars represent standard deviation across triplicate biological samples.

### Growth of C260V mutant and WT *Synechocystis* strains under high light and iron stress

To elucidate the effect of the single amino acid change on cell growth, we compared the growth rates of the C260V mutant and WT *Synechocystis* strains under different conditions (Figure 7A and 7B). Although the growth rates of the two strains were not significantly different from each other, differences in their growth patterns were evident. The initial lag phase was missing in the mutant strain and consequently it grew faster during this phase. Under high light and iron-replete condition, the mutant strain grew significantly faster and reached higher optical density at 730 nm (OD_730_) in three days compared to the WT strain (Figure 7A). This suggested that with sufficient iron in the growth medium, the lack of IsiA-mediated excitation quenching helped accelerate the growth of the mutant cells. Next, to remove trace amounts of iron remaining in the cells that were used as an inoculum, deferoxamine (DFB), an iron-chelator, was added to create stringent iron deficient conditions. Under such severe iron deficiency and high light conditions, the mutant did not show a lag phase and exhibited an initial faster growth rate compared to the WT strain. However, there was a decline in growth after about 30 h, when the cells also showed a bleached phenotype (Figure 7B). In contrast, under low light, the mutant did not exhibit a lag phase and grew as well as WT without bleaching out. These results showed that the C260V mutant is significantly more light-sensitive than the WT strain under severe iron-limited conditions and demonstrated the significant role that the Cys-mediated quenching mechanism in this protein plays in cellular photoprotection.

**Figure 7.**
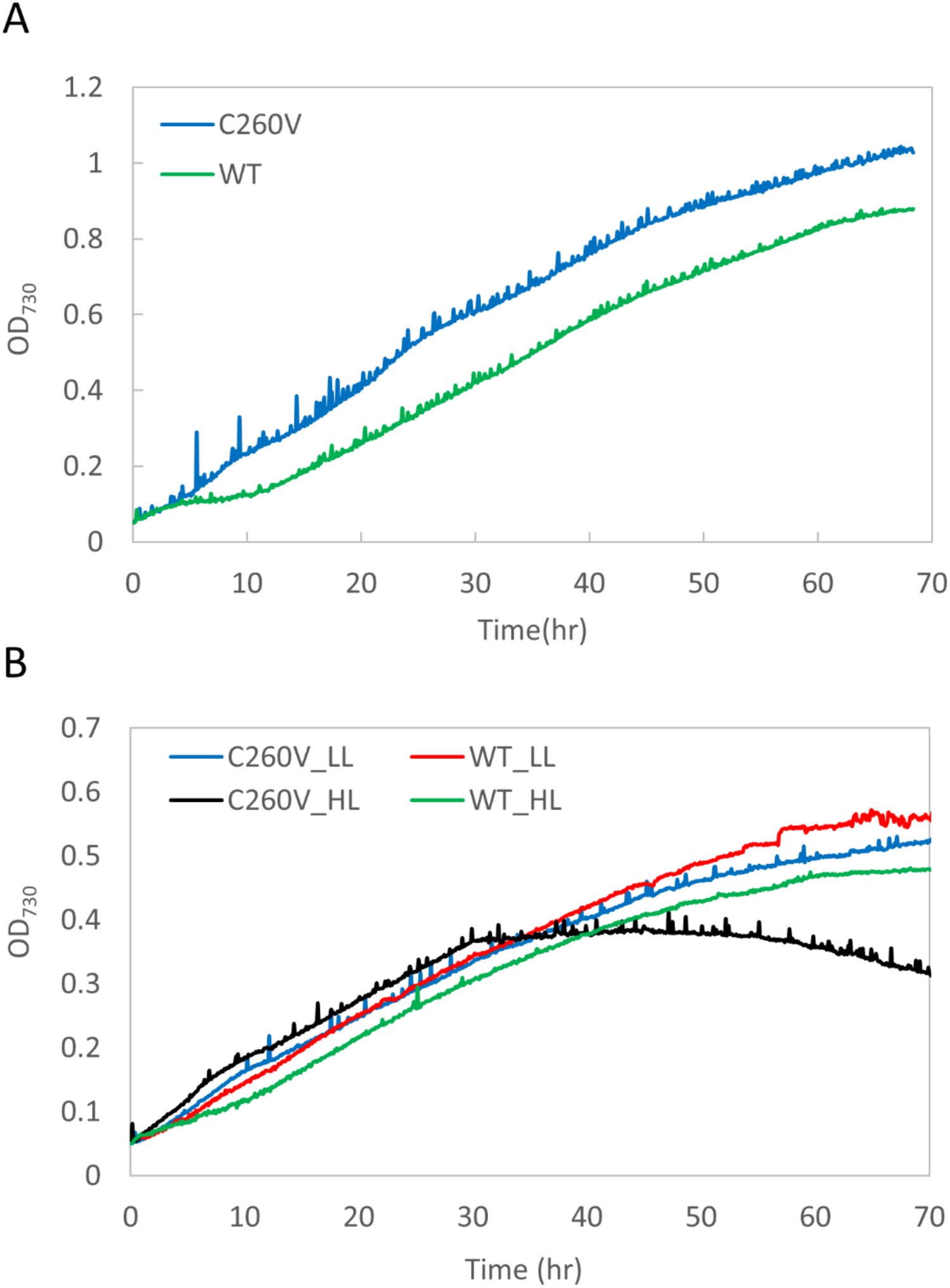
Comparison of growth patterns of WT and C260V strains. Growth curves of C260V and WT cultures under (A) iron-replete conditions under 800 μmol photons m^−2^ s^−1^ (high light, HL) and (B) iron-depleted conditions (with the addition of iron-chelator, DFB) under low light (LL) and high light.

## Discussion

Excessive high light intensities can lead to damages in the cellular machinery of cyanobacteria, algae and plants ^41^. Various mechanisms have evolved in cyanobacteria to avoid such light stress. One such process, unique to these prokaryotes, is based on an orange carotenoid protein (OCP) that can directly quench the excess excitation in the phycobilisomes. The mechanism of this process has been carefully dissected during recent years ^42,43^. Here, we have described another novel and potentially more universal mechanism of photoprotection based on excitation quenching mediated by a Cys residue in the Chl antenna protein IsiA.

### Energy transfer in the mutant C260V IsiA protein

There are three proposed physiological roles of IsiA in cyanobacterial cells. First, in a PSI-IsiA complex, IsiA acts as an efficient accessory antenna for PSI ^21,22,25,44^. Second, IsiA plays a significant role in dissipating excess light energy and thus acts as a cellular photoprotectant ^13,26-^ _29,45_. Third, under stress conditions, IsiA acts as a Chl reservoir and becomes an immediate source of chlorophylls when such stress is relieved and new photosystem complexes are formed ^46-48 49^. In any case, the IsiA only pigment protein complex needs to have an energy dissipation system to avoid deleterious effects of overexcitation and harmful radical formation. In this context, carotenoids in IsiA are in proximity to some Chl *a* molecules^20^, and could likely play a role in excitation energy quenching as previously proposed^31^. However, in a recent study, we have conclusively demonstrated that excitation energy transfer between Chl *a* and carotenoids in IsiA does not occur^32^. Thus, there is a need for an alternate mechanism for quenching of excitation energy in the IsiA complex.

Based on chemical redox titration experiments, we have earlier suggested that IsiA uses a cysteine-mediated mechanism, similar to that in an FMO complex to quench excitation energy ^32,33^. It has been proposed that in FMO under oxidizing conditions, quenching of excitation energy in bacteriochlorophyll (BChl) *a* is facilitated by electron transfer between the excited BChl *a* and the thiyl radical at a cysteine residue ^33^. The rate of photosynthesis is thereby reduced, protecting the photosynthetic apparatus from photodamage. On the other hand, under reducing conditions, the thiyl radical is converted to a thiol group (or thiolate), and, therefore, no excitation quenching takes place.

In contrast to the FMO protein, IsiA has only one cysteine, making it even more crucial in such a quenching process. In this study we performed site-directed mutagenesis to replace the unique cysteine (C260) in IsiA with a valine. The essentially identical absorption spectra of C260V and WT IsiA (as well as WT PSI-IsiA and PSI-C260V IsiA) demonstrate that the C260V IsiA maintains all of the Chl *a* binding pockets with their Chl *a* molecules, suggesting that the C260V IsiA is properly folded. In addition, the Chl *a* Q_y_ absorption bands of both WT and C260V IsiA have the maximum at 670.8 nm, and that of both WT and mutant PSI-IsiA have the maximum at 673.8 nm (Figure 1 C and 1D), in good agreement with previous studies ^18,50^. While subtle structural changes might have occurred in the C260V IsiA protein, these results suggest that we successfully obtained well-folded free C260V IsiA and PSI-C260V IsiA protein in the mutant strain.

According to the recent description of the molecular structure of IsiA in *Synechocystis* 6803 (PDB: 6NWA) at a 3.5 Å resolution^20^, the thiol group of C260 (chain q) lies close to the conserved Chl a5 (505.q), Chl a6 (506.q) and Chl a14 (514.q). In particular, the edge-to-edge distances from Chl a5 and Chl *a*6 are 5.7 Å and 3.7 Å, respectively. These distances are short enough to facilitate electron transfer between these Chl *a* molecules and C260, the basis for the proposed Cys-mediated excitation quenching mechanism^33^. The more recent and higher resolution structures of IsiA from *Synechococcus* 7942 (PDB:6KIF) and *Thermosynechococcus vulcanus* (PDB:6K33) also exhibit similar associations between these conserved chlorophylls and the unique Cys residue^35,36^.

Cyanobacteria are oxygenic organisms and unlike FMO, under physiological conditions, IsiA is expected to be in an oxidizing environment. We have earlier reported that the Chl *a* fluorescence lifetime of IsiA is extended with the addition of a reducing agent^32^ owing to the conversion of its thiyl radical to a thiol group under reducing conditions. This prevents transfer of excited electron from Chl *a* to the thiyl radical, thus decreasing quenching^33^. Our current study shows that the fluorescence decay lifetime of Chl *a* in the C260V IsiA is even longer than that of the WT IsiA under reducing conditions (Figure 3), conclusively demonstrating that quenching in IsiA is critically dependent on C260.

In the PSI-IsiA supercomplex, IsiA functions as an accessory antenna that absorbs light energy and transfers excitation to the reaction center of PSI. Our results are consistent with previous studies, showing that the energy transfer from IsiA to PSI is rapid and efficient^21,22,25^. Moreover, when excited at 660 nm, the mutant PSI-C260V IsiA and the WT PSI-IsiA have identical fluorescence decay traces at 684 nm and 720 nm (Figure 4), indicating identical excitation energy transfer process in both samples. These findings showed that the mutant C260V IsiA was still capable of transferring excitation energy to PSI and served as an accessory antenna for PSI.

### Physiological consequences of the C260V modification

Previous studies showed that IsiA is essential for the survival of *Synechocystis* 6803 and *Synechococcus* sp. PCC 7942 under iron-deficient conditions and under high light^13,27,44,51^. It has been suggested that these cells cannot survive without IsiA mainly due to the photodamage caused in its absence^13,27,44,51^. Our spectroscopic data showed that the mutant C260V IsiA no longer quenches excitation energy (Figure 3) but still functions as an efficient light-harvesting antenna for PSI (Figure 4). We then determined how this single amino acid substitution affects the physiology of the mutant cells.

### Under iron-replete conditions

As discussed earlier, availability of iron in the cells has a profound effect on the expression of the *isiA* gene. When grown under low light with sufficient iron, the absorption spectra of the C260V mutant and the WT cells were almost identical (Figure 5A), and IsiA was absent in either culture (Figure 5B). Under high light, even with sufficient iron, IsiA was induced, as confirmed by the spectroscopic (Figure 5) and immunoblotting (Figure 6C) analysis of the WT and the mutant cells. In fact, the increase in the Chl *a* content in the WT under high light was due to the significant expression of IsiA (Figure 6A and 6C). In contrast, the C260V mutant showed only a slight increase in the Chl *a* content under the same condition, because of its much lower IsiA content. Given that the C260V IsiA cannot quench excitation energy, it is likely that the lower IsiA content in the mutant strain under high light is caused by photodamage.

The growth rates of both strains under low light were nearly identical, but distinct differences in growth patterns were observed. In contrast to the WT, the C260V mutant cells exhibited an initial faster growth rate immediately after inoculation and no lag phase was observed as typically seen for the WT (Figure 7A). Further, under high light conditions, the C260V mutant grew faster than the WT (Figure 7B). Appropriate manipulation of photoprotection has been considered to be one of the attractive approaches to improve photosynthetic yield^52^. It has been shown that by accelerating the recovery from photoprotection or alleviating various photoprotective mechanisms, the growth yields of plants and algae can be substantially improved ^53-55^. As discussed earlier, the mutant C260V IsiA could serve as an efficient light-harvesting antenna for PSI without any quenching and thus improve cell growth under iron replete and high light conditions.

### Under iron-depleted conditions

In the absence of iron, both the WT and mutant strains showed a blue shift of the Chl *a* Q_y_ absorption band, indicating the presence of IsiA under both low and high light conditions (Figure 5C). Under high light, there was a significant reduction in the Chl *a* Q_y_ absorption band in the C260V mutant, indicating a lower Chl *a* content in these cells. As discussed earlier, a proposed function of IsiA is to maintain the cellular Chl *a* content in iron-deficient environments and help the cells recover once iron becomes available^46-48,56^. Our data show that, compared to the C260V mutant, WT cells are better equipped to maintain their cellular Chl *a* content under high light. Furthermore, under high light the photoactive PSI and IsiA contents in the C260V mutant strain were significantly lower (Figure 6B). Because the C260V IsiA is unable to quench excitation energy, it is likely that the loss of PSI and IsiA under high light is due to severe photodamage in this mutant strain. In fact, in a strictly iron-free medium, under high light, the C260V mutant exhibited initial fast growth but then bleached out (Figure 7B). Evidently the C260V mutant was more light-sensitive under iron-depleted conditions and a fully functional IsiA is necessary for the cells to survive high light in iron-depleted environments.

In summary, we have demonstrated that the C260V mutation abolishes the excitation energy quenching ability of IsiA, confirming the critical role of this unique cysteine residue in the quenching process. Our results further showed that when grown under stringent iron deficiency and high light, the mutant strain is more light-sensitive, demonstrating that the cysteine residue in IsiA is crucial for the survival of the cyanobacterial cells in such extreme environments. We also determined that the C260V IsiA serves as an efficient light-harvesting antenna for PSI. Faster growth was observed in the C260V mutant when grown in the presence of iron under high light. This suggests that the single amino acid change may not interfere with other IsiA functions and in fact, light energy utilization may become more efficient in the mutant cells due to the removal of the relevant energy quenching process. Elucidation of this novel Cys-mediated photoprotective quenching mechanism in an oxygenic photosynthetic organism raises the intriguing possibility of occurrence of similar mechanisms in plants and algae. Our findings also provide the framework of engineering such an energy dissipation process in other natural as well as artificial Chl-antenna proteins to modulate photosynthetic productivities under diverse environmental conditions.

## Methods

### Mutant construction

To generate the C260V strain, the mutation was introduced with the CRISPR/Cas12a (Cpf1) system reported previously ^38^. The editing plasmid was constructed by cloning the annealed oligos, the gRNA targeting the *isiA* sequence, into the *Aar*I site on the pSL2680 vector. The repair template was constructed by Gibson assembly to clone two 900 bps homology regions, including the mutation at the PAM sequence and the cysteine coding sequence, into the *Kpn*I site on the editing vector. The resulting plasmid, pSL2854, was verified by sequencing, and transferred to the WT *Synechocystis* cells using the *E. coli* strain containing the pRL443 and pRL623 plasmid in triparental conjugation ^57^. The resulting colonies were repatched three times onto BG11 plates containing 10 μg/mL kanamycin. Mutations were verified by sequencing. The verified colonies were grown to stationary phase in BG11 without antibiotics, diluted 1000 times and grown to stationary phase again. This process was repeated several times to cure the editing plasmid. BG11 plates with and without kanamycin were used to screen the kanamycin-sensitive colonies, which had lost the editing plasmid. Such a kanamycin-sensitive strain was used as the markerless C260V mutant.

A plasmid to generate the C260V-His strain was constructed by replacing the kanamycin resistance gene in the plasmid that was used to generate a IsiA-His strain ^32^ with a gentamicin resistance gene and introducing the site-specific mutation in one of the homologous arms. This plasmid was constructed by Gibson assembly ^58^, using the DNA fragments amplified by PCR. The resulting plasmid pSL2973 was verified by sequencing. The IsiA-His strain was transformed and the transformants were selected for growth on gentamicin. Segregation of the C260V-His strain was confirmed by PCR.

### Culture growth conditions and thylakoid membrane preparation

Wild type and C260V *Synechocystis* cells were grown photoautotrophically in BG11 under 30 μmol photons m^−2^ s^−1^ of white light at 30 °C. After 5 days, cells were harvested and washed three times with YBG11-Fe, a modified medium without any added iron ^32,59^. The washed cells were adjusted to the same optical density of 0.05 at 730 nm and grown under 200 (low light) or 800 (high light) μmol photons m^−2^ s^−1^. For iron-starved liquid cultures, BG11 was replaced with YBG11-Fe with or without the addition of the chelator deferoxamine (DFB) to a final concentration of 50 µM, depending on the experimental settings. The OD_730_ was continuously recorded every 10 minutes over the course of the growth experiments. After three days of growth, cells were harvested and counted. One milliliter of the each culture was used to obtain the absorption spectra, and the rest of the cultures were divided based on the same cell number and resuspended in RB (50 mM morpholineethanesulfonic acid [MES]–NaOH [pH 6.0], 10 mM MgCl_2_, 5 mM CaCl_2_, 25% glycerol) and then stored at −80 °C for future use.

The cells were thawed on ice prior to the thylakoid membrane extraction. Cells were broken by bead-beating as described previously ^60,61^ with following modifications. The thawed cells and 0.17 mm glass beads were loaded into a prechilled Eppendorf tube in a 1:1 ratio of cell suspension to glass beads. Cells were then broken using 10 break cycles, each cycle consisting of 1 min of homogenization on a Vortex mixer, followed by 1 min of cooling. Cell homogenates were centrifuged at 30,000 *×* g for 15 min and washed with RB once. The resulting pellet was resuspended in RB and solubilized with β-D-dodecyl maltoside (DDM) (final concentration of 1%), and incubated on ice in dark with gentle stirring for 30 min. The sample was then centrifuged at 30,000 *×* g for 30 min and the resulting supernatant, the solubilized thylakoid membranes, was stored at −80 °C for future use.

### Cell counting

Cell cultures were grown in MC-1000 multicultivators in BG11 and YBG11-Fe under 200 or 800 μmol photons m^−2^ s^−1^ as mentioned above. The cells were harvested after three days, diluted to OD_730_ = 0.01 and the cell number was counted on an automated cell counter (Cellometer Vision; Nexcelom) with further manual curation to improve accuracy.

### Photoactive PSI content

Cells were harvested after three days and adjusted to same cell numbers. With the addition of 10 μM 3-(3,4-dichlorophenyl)-1,1-dimethylurea (DCMU) and 20 μM dibromothymoquinone (DBMIB), which block linear and cyclic electron flow respectively, the absorbance at 705 nm of P700 in each sample was recorded for 5 s under saturating light on a JTS-10 pump probe spectrophotometer (Bio-Logic Instruments). A molar extinction coefficient of 70 mM^-1^ cm^-1^ for P700 was used to estimate the photoactive PSI content.

### SDS-PAGE and immunoblot analysis

Solubilized thylakoid membranes of C260V and WT strains were prepared as mentioned above. The solubilized thylakoid membranes were fractionated by denaturing sodium dodecyl sulfate polyacrylamide gel electrophoresis (SDS-PAGE) and analyzed by protein immunoblot as described earlier ^62,63^. Fractionated proteins were blotted onto polyvinylidene difluoride (PVDF) membranes. IsiA and D2 were identified by using specific antisera and visualized by using enhanced chemiluminescence reagents (WestPico; Pierce) on an Odyssey Fc imager (LI-COR Biosciences, USA). The relative protein content was estimated by Image Studio (LI-COR Biosciences, USA) based on the chemiluminescence signals of samples.

### Pigment content estimation

Chl *a* concentration was estimated by a methanol extraction method^64^. To determine phycobilin concentration, absorption spectra of cultures were obtained on a DW2000 spectrophotometer (OLIS, USA), using the following equation^65-67^:

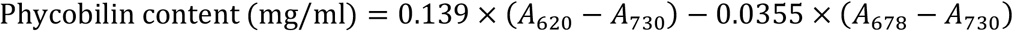

*A*_620_, *A*_678_ and *A*_730_ are absorbances at 620, 678 and 730 nm, respectively, in a 1 cm path length.

### Protein complex purification

C260V-His liquid culture was grown in YBG11-Fe with the addition of 5 mM deferoxamine under 30 μmol photons m^−2^ s^−1^ at 30 °C for two weeks to induce *isiA* expression as described earlier ^32^. The solubilized thylakoid membranes were prepared as described above, and then used to purify PSI-IsiA and IsiA protein complexes, using nickel affinity chromatography and rate-zonal centrifugation ^32^. After 18 h of ultracentrifugation, fractions were collected. The first and fourth green bands from the top of the gradient contained IsiA and PSI-IsiA supercomplexes, respectively (Fig. 2), as determined by immunoblot and spectroscopic analysis.

### Steady-state and time-resolved spectroscopy

Steady-state absorption spectra were recorded using a UV-1800 spectrophotometer (Shimadzu). Time-resolved fluorescence (TRF) experiments were carried out using two different setups. For recording of image of fluorescence profiles of PSI-IsiA supercomplexes at 77 K, a setup based on Hamamatsu (Japan) universal streak camera described previously^68^ was used. Single wavelength traces for fluorescence decay of IsiA samples at room temperature were recorded using a standalone Simple-Tau 130 time-correlated single photon counting (TCSPC) system from Becker & Hickl (Germany). Both setups were coupled to an ultrafast laser system (Spectra-Physics, USA) described previously^69^. The frequency of the excitation pulses was set to 8 MHz, corresponding to ∼120 ns between subsequent pulses. To minimize the detection of scattered light from the excitation beam, a long-pass 665 nm filter was placed at the entrance slit of the spectrograph/monochromator. The integrity of the samples was examined by monitoring the realtime photon count rate over the time course of the experiment. The samples were resuspended to an absorbance of ≤0.1 at the Chl *a* Q_y_ band and the emission signal was recorded at a right angle with respect to the excitation beam. The excitation beam set to 640 nm, with photon intensity of ∼10^10^ photons/cm^2^ per pulse was depolarized and focused on the sample in a circular spot of ∼1 mm diameter. Fluorescence emission was recorded at 684 nm.

## Acknowledgements

This study was supported by the Chemical Sciences, Geosciences, and Biosciences Division, Office of Basic Energy Sciences, Office of Science, US Department of Energy (DOE) Grant DE-FG02-99ER20350 (to H.B.P.). We thank all members of the research group of Himadri Pakrasi for critical scientific discussions. H.-Y.S.C., D.M.N. and H.B.P designed the experiments; H.-Y.S.C., D.M.N. A.B. and S.B. performed the experiments; H.-Y.S.C., D.M.N., A.B. and H.B.P. wrote the paper.

## Conflict of Interest

The authors declare no conflict of interest.

## Notes

### Competing Interest Statement

The authors have declared no competing interest.

